# Effectiveness and efficiency: label-aware hierarchical subgraph learning for protein-protein interaction

**DOI:** 10.1101/2024.03.08.584199

**Authors:** Yuanqing Zhou, Haitao Lin, Yufei Huang, Lirong Wu, Stan Z. Li, Wei Chen

**Author notes:** **Corresponding authors:** Prof. Wei Chen, Department of Food Science and Nutrition, College of Biosystems Engineering and Food Science, Zhejiang University, Hangzhou, 310058 China., Prof. Stan Z. Li, School of Engineering, Westlake University, Hangzhou, 310024 China.

## Abstract

Protein-protein interactions (PPIs) are crucial in various biological processes and their study has significant implications for drug development and disease diagnosis. Existing deep learning methods for PPIs prediction, including graph neural networks (GNNs), have been widely employed as the solutions, while they usually suffer from performance degradation under complex real-world scenarios. We claim that the topological shortcut is one of the key problems contributing negatively to the performance, according to our analysis. By modeling the PPIs as a graph with protein as nodes and interactions as edge types, the prevailing models tend to learn the pattern of nodes’ degrees rather than intrinsic sequence-structure profiles, leading to the problem termed topological shortcut. In addition, with the emergence of high-throughput experimental methods such as mass spectrometry and protein chip technology, the amount of available PPI data is exploding. The huge data growth leads to intensive computational costs and challenges computing devices, causing infeasibility in practice. To address the discussed problems, we propose a **l**abel-**a**ware hie**r**archical s**u**b**g**raph **l**earning method (laruGL-PPI) that can effectively infer PPIs while being both interpretable and generalizable. Specifically, we introduced edge-based subgraph sampling to effectively alleviate the problems of topological shortcuts and high computing costs. Besides, the inner-outer connections of PPIs are modeled as a hierarchical graph, together with the dependencies between interaction types constructed by a label graph. Extensive experiments on PPI datasets of different scales demonstrate that laruGL-PPI outperforms state-of-the-art PPI prediction methods, particularly in the testing of unseen proteins. Also, our model can recognize crucial sites of proteins, such as surface sites for binding and active sites for catalysis.

## 1 Introduction

Biological functions are accomplished by interactions and chemical reactions among biomolecules. Among them, protein-protein interactions (PPIs) are undoubtedly one of the most essential molecular events in the human body and are an important source of therapeutic interventions against diseases. A comprehensive dictionary of PPIs can help connect the dots in complicated biological pathways and expedite the development of therapeutic[1, 2]. In past decades, high-throughput experimental methods, e.g. yeast two-hybrid screens (Y2H)[3], and mass spectrometric protein complex identification (MS-PCI)[4] have been developed to identify PPIs. Nevertheless, genome-scale experiments are expensive, tedious, and time-consuming while suffering from high error rates and low coverage[5]. Therefore, it is urgent to establish computational approaches with high accuracy and speed to identify potential PPIs.

Recently, a large variety of computational approaches have been developed to detect PPIs, consisting of classic machine learning (ML)-based [6-8] and deep learning (DL)-based methods[9-12]. Compared with classic ML methods, the DL algorithm has an advantage in processing complicated and large-scale data and extracting useful features automatically, achieving huge success in computational biology[13], including PPI prediction[14]. With the development in graph neural networks (GNNs), more recent works focus on modeling the PPIs as a graph, by constructing a PPIs graph with proteins as nodes and interactions as edge types, with various GNNs [15, 16] utilized to explore the PPIs graph to facilitate inference of type of interactions[12, 17, 18]. Among these graph-based methods, the HIGH-PPI reaches state-of-the-art performance with interpretability. However, HIGH-PPI suffers from intensive computational costs due to hierarchical graphs with protein structures and PPI networks.

More importantly, we find out that topological shortcut (TS)[19] exists in multiple types PPI prediction. The emergence of TS causes DL-based methods difficult to learn from protein. The TS refers to the fact that, in the PPI network, the model easily learns simple topological patterns during the training process, without fully exploiting the 1D protein sequences, 3D structures, and physicochemical properties profiles closely related to PPI. As a result, in the inference process, the model makes use of known topological knowledge to predict PPI, while seldom considering the characteristics of the protein itself. This makes the model collapse when faced with ‘unseen’ proteins. Experimental results demonstrate the unsatisfactory performance in inductive and BFS scenarios for the previous models (See section 4).

In this work, to tackle the above problems, we proposed an interpretable and generalizable PPI prediction framework with edge-based subgraph sampling, referred to as **l**abel-**a**ware hie**r**archical s**u**b**g**raph **l**earning framework (laruGL-PPI). Firstly, we propose leveraging graph structure to model protein relations and label dependencies for multi-graph learning. In detail, we derive an inter-dependent classifier to extract information from the label graph, which is then employed to the protein representations aggregated by neighbors in the PPI network for multi-type PPI prediction. Secondly, we also exploit graphs to depict protein structure, represented with nodes as residues and edges as contacts between them. Thirdly, we adopt edge-based subgraph sampling to alleviate the topological shortcut and the computational cost. To the best of our knowledge, this is the first study to investigate the emergence of topological shortcut and to mitigate it in PPI predictive tasks. Precisely, the main contributions of the work can be summarized as follows:

- For multi-type PPI prediction, we first investigate topological shortcut, which limits learning ability of model. An effective **l**abel-**a**ware hie**r**archical s**u**b**g**raph **l**earning-based PPI prediction (laruGL-PPI) framework was proposed to address this problem.
- In laruGL-PPI, Multiple graphs were constructed to model protein structure, connections between proteins and label dependencies simultaneously. Subgraph sampling was further adopted to train model for enhancing generalization ability of model and reducing computational pressure on devices.
- Extensive experiments on three PPI datasets with different settings demonstrate that laruGL-PPI outperforms other state-of-the-art methods for multi-label PPI prediction under various challenging scenarios and has a high interpretability for PPI type prediction.

## 2. Problem statement and related work

### Protein-Protein Interaction Prediction

Early works leverage machine learning (ML) techniques to map pairs of handcrafted sequence features of proteins to interaction patterns[6, 8, 20, 21]. With the development of deep learning (DL), recent works have applied deep neural networks to automatically extract features from protein sequences for enhancing feature representation[10, 11, 22]. Furthermore, the latest works utilize graph neural networks (GNN) to model the PPI network[12]. Among these works, the SemiGNN-PPI[17] establishes a multi-graph learning framework to improve the model performance, and the HIGH-PPI[18] organizes the structure-based internal graph for each protein to enhance the interpretability of the model. However, all the PPI prediction models face a topological shortcut problem due to annotation imbalance. Also, the framework using a hierarchical graph with protein structure exerts pressure on computational devices because of the huge memory cost.

### Topological Shortcut

In drug-target interaction (DTI) prediction, similar to PPI, DL models learn a map from features to interaction labels in the training data. Features refer to the chemical structures of proteins and ligands, which determine their physical and chemical properties. In an ideal scenario, the ML model learns the patterns characterizing the features that drive the protein-ligand interactions, capturing the physical and chemical properties of a protein and of a ligand that determine the interaction. Nevertheless, multiple state-of-the-art deep learning models ignore the features and depend largely on annotations, i.e., the degree information for each protein and ligand in the drug-target interaction (DTI) network, as a shortcut to make new binding inferences. In protein-protein interaction prediction, the topological shortcut problem also exists in the PPI network.

### Multi-Label Learning

MLL addresses the problem in assigning multiple labels to a single instance. It has been applied successfully in various domains, e.g., computer vision[23, 24].

Traditional MLL methods typically train independent classifiers for all labels but fail to pay attention to the potential label interdependence, leading to suboptimal performance. Recent trends in MLL incorporate deep learning to capture the label dependencies[25, 26]. As an example, CNN-RNN[25] makes use of recurrent neural networks (RNNs) to transform the label vectors into an embedded space to learn label correlations implicitly. More recently, graph-based MLL approaches have gotten great attention from scientific communities[27, 28]. Especially, the SemiGNN-PPI[17] successfully makes use of Graph Convolutional Network (GCN) by constructing a directed graph over labels to explicitly model the protein-protein interaction dependencies adaptively. In this regard, we add graph-based MLL to hierarchical graph learning for more accurate PPI prediction.

### Subgraph Sampling

In a GCN, data to be gathered for one output node comes from its neighbors in the previous layer. Each of these neighbors, in turn, gathers its output from the previous layer, and so on. Clearly, the deeper we back track, the more multi-hop neighbors to support the computation of the root. The number of support nodes (and thus the training time) potentially grows exponentially with the GCN depth. To mitigate such “neighbor explosion”, state-of-the-art methods use various layer sampling techniques[29-31]. While these methods significantly speed up training, they face challenges in scalability, accuracy, or computation complexity. Recently, the GraphSAINT[32] was presented to efficiently train deep GCNs. Instead of building a GCN on the full training graph and then sampling across the layers, GraphSAINT samples the training graph first and then builds a full GCN on the subgraph, which is graph sampling-based. In this work, we leverage edge-based subgraph sampling to alleviate the topological shortcut problem in the PPI network and computational cost.

## 3. Materials and methods

### 3.1 Task definition

Given a set of protein *P* = {*p*_0_, *p*_1_,…, *p*_*n*_ }and a set of PPIs *E* = {*e*_*ij*_ = { *p*_*i*_, *p* _*j*_ } | *i* ≠ *j, p*_*i*_, *p* _*j*_ ∈ *P, I* (*e*_*ij*_) ∈ 0,1}, where *I* (*e*_*ij*_) is a binary PPI indicator function that is 1 if the PPI between proteins *p*_*i*_ and *p* _*j*_ has been confirmed, and 0 otherwise, the types of PPI can be represented by the label space *C* ={*c*_1_, *c*_2_,…, *c*_*t*_ }with *t*different types of interactions, and the labels for a confirmed PPI *e*_*ij*_ can be represented as *y*_*ij*_ ⊆ *C* . The goal of multi-type PPI learning is to learn a function 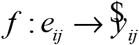 from the training set such that for any PPI 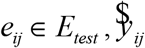 is the set of predicted labels for *e*_*ij*_ . To confirm the emergence of topological shortcuts in PPI prediction, we analyze annotations in three common PPI datasets (i.e., SHS27k, SHS148k, and STRING).

### 3.2 Network architecture design of laruGL-PPI

**Figure 1** depicts the overview of our proposed laruGL-PPI framework. We first construct the multi-graph encoding module to effectively predict PPI types, consisting of a protein graph encoding network for capturing binding patterns, a PPI network graph encoding network for modeling protein-protein correlations, and a label graph encoding network for learning label dependencies. To mitigate topological shortcut problem and computational pressure, we make use of an edge-based subgraph sampling approach to train the laruGL-PPI model.

**Figure 1:**
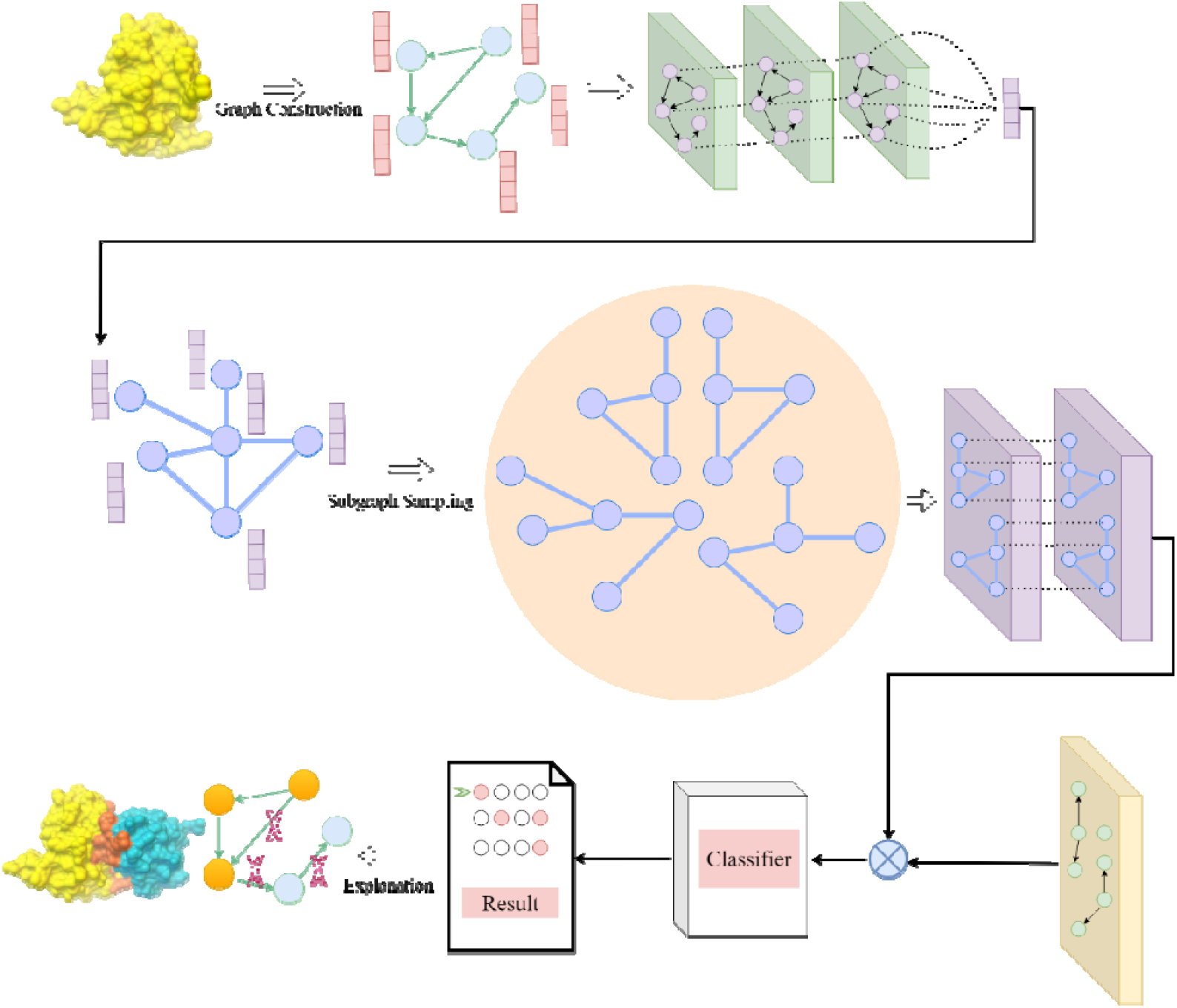
The overall framework of laruGL-PPI. First, we construct a protein graph with residues as nodes and contacts as edges. We treat the physical and chemical properties of amino acids as features of nodes. Then, GCN blocks are employed to characterize the protein graph, and all the output features of nodes are aggregated to a feature vector as a feature of the protein node in the PPI network graph, which is characterized by proteins as nodes and correlations as edges. Afterward, we use GIN blocks to model the PPI network, leading to network-aware protein features. We arrange GCN blocks on the label graph to learn a classifier, which is applied to the products of two protein features for protein-protein interaction prediction. Later on, laruGL-PPI is interpreted by GNNExplainer.

#### 3.2.1 Multi-Graph Encoding

##### Protein-Graph Encoding

We denote a set of amino acid residues in a protein as p*rot* = {*r*_1_, *r*_2_,…, *r*_*n*_ }. Each residue is described with *t*kinds of physicochemical properties. For the bottom inside-of-protein view, a protein graph *g*_*b*_ = (*V*_*b*_, *A*_*b*_, *X*_*b*_) is built to model the relationship between residues in Prot, where *V*_*b*_ ⊆ *prot* is the set of nodes, *A*_*b*_ is an *n*×*n* adjacency matrix representing the connectivity in *g*_*b*_, and *X* ∈ *R*^*n*×*θ*^ is a feature matrix containing the properties of all residues. The contact map is exactly equivalent to the adjacency matrix *A* ∈{0,1}^*n*×*n*^ in the protein graph. Contact maps are obtained with atomic level 3D coordinates of proteins. First, we retrieve the predictive structures from the AlphaFold Protein Structure Database. Then we represent the location of each residue by the 3D coordinates of its *C*_*α*_ atom. The presence or the absence of contact between a pair of residues is determined by their *C*_*α*_ − *C*’_*α*_ physical distance. In this work, we choose the cutoff distance of 12 A. For a feature matrix *X* _b_, each row stands for a set of properties for one amino acid residue. In this context, seven residue-level properties are considered: isoelectric point, polarity, acidity and alkalinity, hydrogen bond acceptor, hydrogen bond donor, octanol-water partition coefficient, and topological polar surface area. We apply variant graph neural networks to learn protein representations. Specifically, four GCN blocks are employed. Given the adjacency matrix *A*_*b*_ ∈{0,1}^*n*×*n*^ and the feature matrix *X*_*b*_∈ *R*^*n*×*θ*^ of an arbitrary protein graph *g*_*b*_, we update residue representations with the neighbor aggregations based on the work of Kipf and Welling:

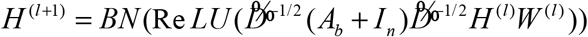

where *I* ∈ *R*^*n*×*n*^ is the identity matrix, 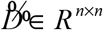 is the diagonal degree matrix with entries 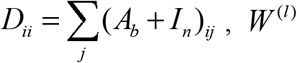 is a learnable weight matrix for the GCN layer *l*, ReLU, and BN denotes the ReLU activation function and batch normalization, respectively. In the first GCN layer, *H* ^(0)^ = *X*_*b*_ . After four rounds of iteration, we perform the readout operation with a self-attention graph pooling layer and average aggregation to obtain the entire graph representation.

##### PPI-Graph Encoding

The PPI network structure determines the adjacency matrix *A* _*t*_ ∈{0,1}^*m*×*m*^, in which the *i* -th row and *j* -th column elements are 1 if the *i* -th protein interacts *j* -th. The *i* -th row of the feature matrix *X*_*t*_ describes the representation vector for the *i* –th protein graph *g*_*b*_ . We use a graph isomorphism network (GIN) to learn PPI network information.

Formally, we are given the PPI graph *g*_*t*_ = (*V*_*t*_, *A*_*t*_, *X*_*t*_), where *X* _*t*_ ∈ *R*^*m*×*d*^ is defined as the feature matrix whose row vector is a representation from protein graph. In the *k* -th GIN layer,

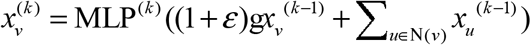

where 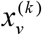 denotes the representation of protein *v*after the *k* -th GIN layer, N(*v*) is a set of proteins adjacent to *v*, and *ε* is a learnable parameter. After two GIN layers, we feed 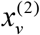 in a linear layer with the ReLU activation to obtain the final protein representation. For an arbitrary query pair containing the *i*-th and *j*-th proteins, we adopt the element-wise product to combine protein pair information. Then the generated vector goes through a fully connected layer (FC) to come by protein-protein representation.

##### Label-Graph Encoding

In multi-label PPI prediction, correlations exist among different types of interactions, i.e., some PPI types may appear together frequently while others rarely appear together. For example, post-translational modifications are often associated with catalysis in biology. Following Chen et al, we model the interdependencies between different PPI types(labels) using a graph and make the model learn an inter-dependent classifier with the GCN, which is directly applied to protein-protein representation for multi-type PPI prediction. The GCN aims to learn a function *f* (·, ·) on the graph with *t* nodes. With the convolutional operation, *f* (·, ·) can be expressed as:

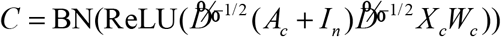

where *I*_*n*_ ∈ *R*^*n*×*n*^ is the identity matrix, 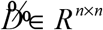 is the diagonal degree matrix with entries 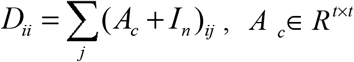and 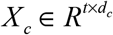 denotes adjacency and feature matrix in label graph individually, *W*_*c*_ is a learnable weight matrix for the GCN, ReLU, BN denotes the ReLU activation function and batch normalization, respectively. Considering that PPI type names are semantic, we utilize the BioWordVec model pre-trained on the biomedical corpus for generating word embedding *X* _*c*_ to better capture their semantics. To construct the label correlation matrix, we compute the conditional probability of different labels within the dataset. To avoid noise, we binarize the correlation matrix with a threshold *τ* to obtain *A*_*c*_ .

#### 3.2.2 Multi-Graph Learning with Subgraph Sampling

By applying the learned classifier *W* from label graph encoding(LGE) to the learned representations from hierarchical graph encoding (HGE) for PPI e_*ij*_, we can obtain the predicted scores 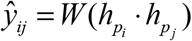. We use the traditional multi-label classification loss function to update the whole network in an end-to-end manner. The loss function can be written as:

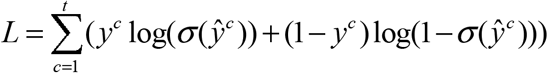

where *σ* (·) is the sigmoid function. Our model learns the aggregated features by combining protein neighbors and models the label correlations by learning inter-dependent classifiers simultaneously to improve the model generalization. With multi-graph learning, the learned classifiers are expected to be neighborhood-aware at both feature and label levels. We select an edge-based sampler to build a subgraph *G*_s_ due to the nature of protein-protein interaction. Indetail, Given training PPI-network graph *G* and edge budget *m* . Firstly, sampling probability *P*((*u, v*)) is calculated by the following formula:

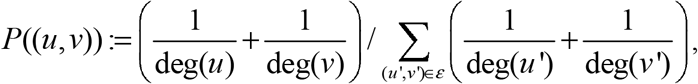

where deg(·), *ε* are referred to as the degree of a protein node and a set of all edges in the training graph. Secondly, *m*edges are randomly sampled from *ε* according to *P* and sets of nodes that are end-points of edges in *ε*_*s*_ are determined as*V*_*s*_ . Finally, the subgraph *G*_*s*_ is derived from*V*_*s*_ with node inducing.

#### 3.2.3 Residue importance computation

We employ the method called GNNExplainer[33] to generate explanations for laruGL-PPI. By taking the well-trained GNN model and its predictions as inputs, GNNExplainer returns the most important subgraph by maximizing the mutual information *MI* between the model prediction and possible subgraphs. Given protein graphs *G*_1_ and *G*_2_ that connect in the PPI network, our goal is to identify a subgraph *G*_s_ ⊆ *G*_1_ that is important for our model’s prediction *ŷ* . The mutual information *MI* is formulated as

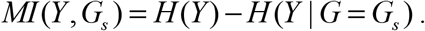

*MI*_*s*_ represents importance of *G*_*s*_, *Y* is a variable indicating the probability of PPI presence of *G*_1_ and *G*_2_, and *H* (g) is the entropy term. In effect, *MI* quantifies the change in the probability of prediction *ŷ* = Φ(*G*_1_, *G*_2_) when the computation graph is limited to explanation subgraph *G*_*s*_ . A computationally efficient version of GNNExplainer’s objective, which we optimize using Adam, is as follow:

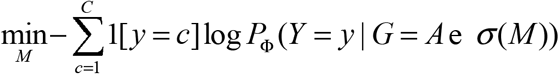

Lastly, we compute the element-wise multiplication of *σ* (*M*) and adjacency matrix *A* and set a threshold to arrive at the explanation *G*_*s*_ for our model’s prediction *ŷ* at the protein graph *G*_1_ .

#### 3.2.4 Datasets and metrics

We perform extensive analyses and experiments on three benchmark datasets, i.e., SHS27k, SHS148k, and STRING. Both SHS27k and SHS148k are subsets of the STRING datasets. These are multi-label PPI data of Homo sapiens from the STRING database. We retrieve protein structures from the AlphaFold Protein Structure Database. In this work, SHS27k has 1553 proteins and 6660 protein-protein multi-type labels. SHS148k consists of 4072 proteins and 30067 labels. The PPIs are annotated with 7 types, i.e., Activation, Binding, Catalysis, Expression, Inhibition, Post-translational modification (Ptmod), and Reaction. Each PPI is labeled with at least one of them. The micro-F1 score was applied to evaluate and compare the predictive performance of laruGL-PPI and existing methods. We also use the area under the precision-recall curve (AUPR) to estimate the model with regard to specific interaction type.

## 4 Results and discussion

### 4.1 Topological Shortcut Analysis

In the context of the PPI predictive task, ML models learn a map from features to labels as annotations and then make inferences for the likelihood of each node (protein) to bind to other nodes. Annotations capture protein-protein interactions. Normally, known protein-protein interaction is denoted as 1, while the unknown interaction is labeled as 0. Features refer to the chemical structures of proteins, which determine their physical and chemical properties, and are expressed as amino acid sequences or 3D structures for proteins. With the success of network inference in various fields, many ML models have considered PPI network and regard it as one of the features externally. In an ideal scenario, the ML model learns the patterns characterizing the features that drive the protein-protein interactions, capturing the physical and chemical properties of protein pairs that determine the mutual binding action. However, as we show next, multiple state-of-the-art deep learning models, such as HIGH-PPI rely largely on the degree information for each protein in the protein-protein interaction network, as a shortcut to make new binding predictions. We begin by noticing that the number of annotations linked to a protein follows a fat-tailed distribution, indicating that the vast majority of proteins have only a small number of annotations, which then coexist with a few hubs, nodes with an exceptionally large number of binding records.

Distributions of the number of annotations in the SHS27k and SHS148k are shown in **Figure 2**. As an example, the number of annotations for proteins follows a power law distribution with degree exponent *γ* = 1.46, maximum degree *k*_max_ = 182, and minimum degree *k*_min_ = 1 on the SHS27k dataset. For the degree exponent, the second moment of the distribution diverges, implying that the expected uncertainty in the binding information is highly significant, limiting our ability to predict the binding between proteins in terms of intrinsic protein characteristics. Furthermore, it is found that the number of annotations *k* and the average interaction per *k*, calculated as the average across all links stemming from nodes of degree *k*, are associated with each other. In this work, there are seven interaction types between proteins, consisting of reaction, binding, posttranslational modification, activation, inhibition, catalysis, and expression. For catalysis, they are positively correlated with *r*_*Spearman*_ = 0.37 (*p* < 0.01), indicating a stronger catalysis propensity for proteins with more annotations. Whereas, they are negatively correlated with *r*_*Spearman*_ = −0.34 (*p* < 0.01) for expression, implying a stronger expression tendency for proteins with fewer annotations (Table 1). As the annotations follow fat-tailed distributions, the observed correlation drives the hub proteins to have disproportionately more interaction or non-interaction on average, whereas proteins with fewer annotations have both interaction and non-interaction examples. This annotation imbalance prompts the ML models to leverage degree information (positive and negative annotations) in making interaction predictions instead of learning interaction patterns from the protein intrinsic structures.

**Table 1.**
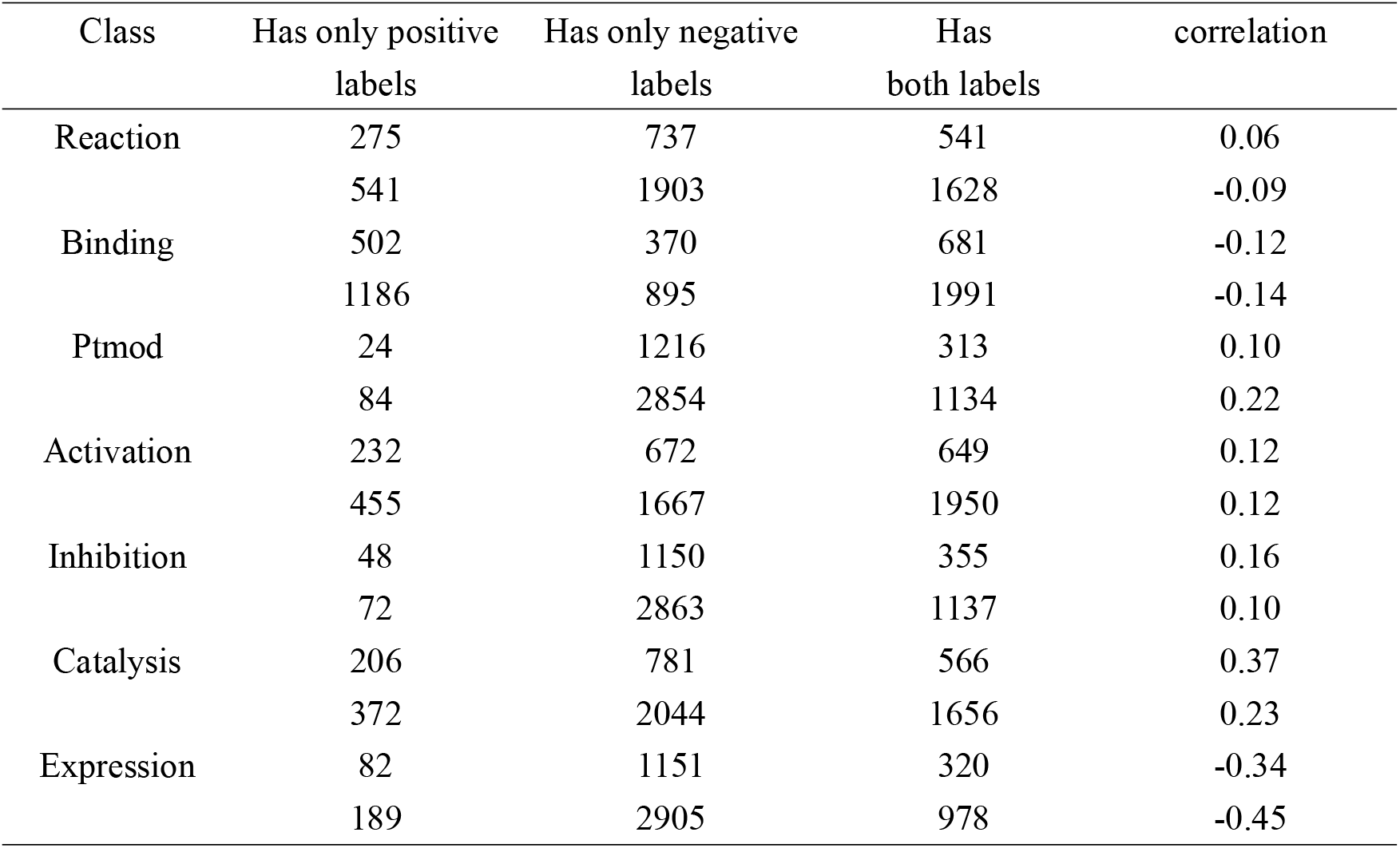
Statics of SHS27K and SHS148K. Most proteins in common datasets have either positive or negative labels, which creates an imbalance in the degree ratio.

| Class | Has only positive<br>labels | Has only negative<br>labels | Has<br>both labels | correlation |
| --- | --- | --- | --- | --- |
| Reaction | 275 | 737 | 541 | 0.06 |
|  | 541 | 1903 | 1628 | -0.09 |
| Binding | 502 | 370 | 681 | -0.12 |
|  | 1186 | 895 | 1991 | -0.14 |
| Ptmod | 24 | 1216 | 313 | 0.10 |
|  | 84 | 2854 | 1134 | 0.22 |
| Activation | 232 | 672 | 649 | 0.12 |
|  | 455 | 1667 | 1950 | 0.12 |
| Inhibition | 48 | 1150 | 355 | 0.16 |
|  | 72 | 2863 | 1137 | 0.10 |
| Catalysis | 206 | 781 | 566 | 0.37 |
|  | 372 | 2044 | 1656 | 0.23 |
| Expression | 82 | 1151 | 320 | -0.34 |
|  | 189 | 2905 | 978 | -0.45 |

**Figure 2.**
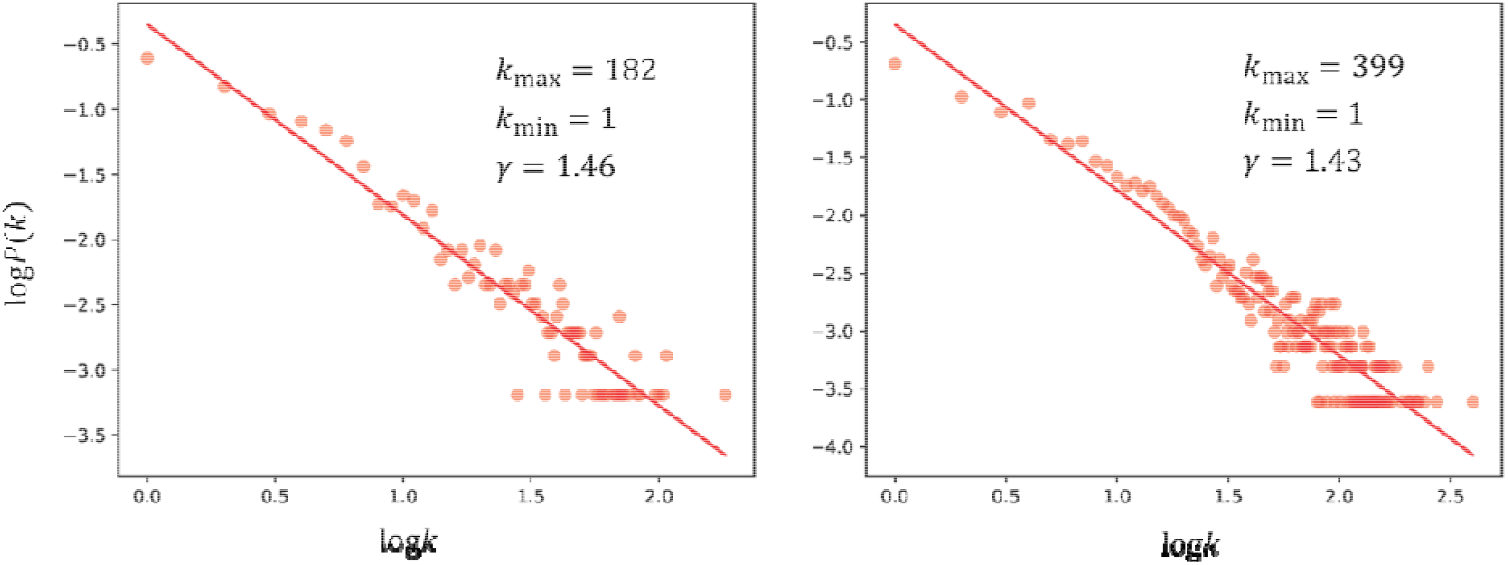
Distributions of the number of annotations in the SHS27k (a) and SHS148k (b)

To investigate the emergence of topological shortcuts, for each node *i* with the number of annotations *k*_*i*_, we quantify the balance of the available data information via the degree ratio, 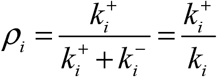, where, 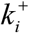 is the positive degree, corresponding to the number of known binding annotations in the training data, and 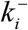 is the negative degree, or the number of unknown annotations in the training data. As most proteins lack either binding or unknown annotations for interaction types (Table 1), the resulting *p*_*i*_ are close to 1 or 0 (See Figure 3), which represents the annotation imbalance in the prediction problem. As many state-of-the-art deep learning models, such as HIGH-PPI, uniformly sample the available positive and negative annotations, they tend to assign a higher binding probability to proteins with higher *ρ* . Consequently, their binding predictions are driven by topological shortcuts in the protein-protein network, which are associated with the positive and negative annotations present in the dataset rather than the structural features characterizing proteins.

**Figure 3.**
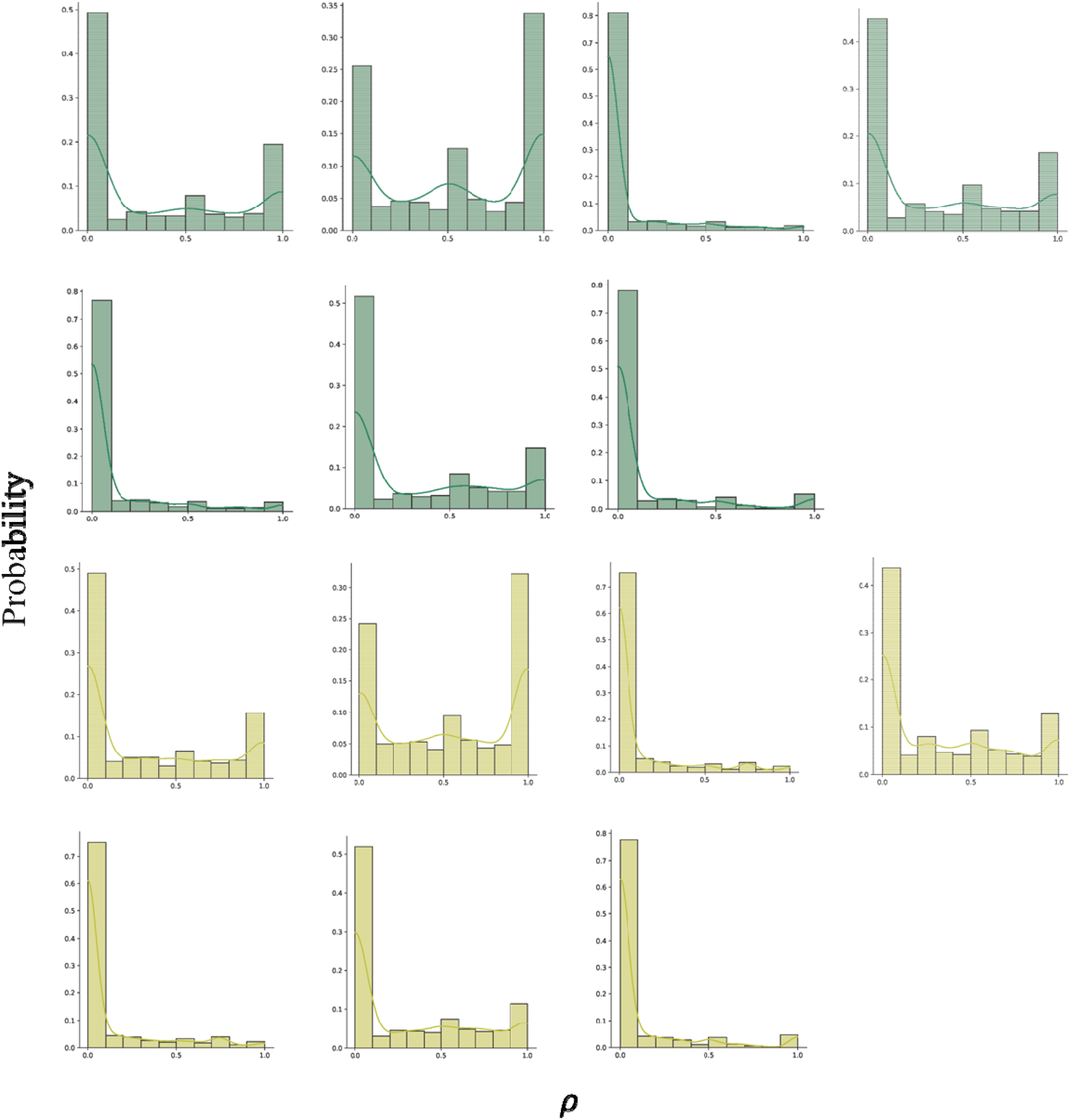
The distribution of degree ratios for the proteins in SHS27K and SHS148K.

### 4.2 Comparison with competitive methods

In this section, we display the performances achieved by our laruGL-PPI and competitive baselines on SHS27k and SHS148k datasets. Here, we ran each model five times independently to eliminate bias caused by randomness and examine the robustness of laruGL-PPI. In Table 2, we compare our methods with other baseline methods under different partition schemes and various datasets. It is observed that graph-based methods, i.e., GNN-PPI, SemiGNN-PPI, and HIGH-PPI exceed other ML and DL models, even under more difficult BFS and DFS partitions with more unseen proteins. It can be ascribed to graph learning, which can model correlations between proteins despite the existence of more unknown proteins. Furthermore, our proposed method, which incorporates multiple graph learning in conjunction with subgraph sampling, achieves competitive performance in multi-type PPI prediction in a common transductive setting. It is worth mentioning that our method reaches state-of-the-art levels in difficult data partition, i.e., DFS and BFS. In the realistic application of biology, models tend not to have seen predicted protein pairs in the training process. To conform with reality, it is the first time to consider an inductive setting in multi-type PPI prediction. When facing unseen protein, the model cannot utilize degree information to conduct prediction, leading to decreasing accuracy. The inductive setting requires models to more focus on protein structures instead of degree information. By subgraph sampling, our model improves predictive performance and outperforms state-of-the-art HIGH-PPI under the more difficult inductive setting with SHS27k, regardless of partition mode (Table 2).

**Table 2.** Prediction accuracy of laruGL-PPI and other competing methods. The accuracy is represented by the mean micro-F1 score (standard deviation). The optimal value in settings has been emphasized in bold. The symbol * stands for inductive setting and the model without * means transductive.

| Model | SHS27K |  |  | SHS148K |  |  |
| --- | --- | --- | --- | --- | --- | --- |
|  | Random | DFS | BFS | Random | DFS | BFS |
| RF | 0.7845 | 0.3555 | 0.3767 | 0.8210 | 0.4326 | 0.3896 |
|  | (0.0088) | (0.0222) | (0.0157) | (0.0020) | (0.0343) | (0.0194) |
| LR | 0.7155 | 0.4851 | 0.4306 | 0.6700 | 0.5109 | 0.4745 |
|  | (0.0093) | (0.0187) | (0.0505) | (0.0007) | (0.0209) | (0.0142) |
| DPPI | 0.7399 | 0.4612 | 0.4143 | 0.7748 | 0.5203 | 0.5212 |
|  | (0.0504) | (0.0302) | (0.0056) | (0.0139) | (0.0118) | (0.0870) |
| DNN-PPI | 0.7789 | 0.5434 | 0.4890 | 0.8849 | 0.5842 | 0.5740 |
|  | (0.0497) | (0.0130) | (0.0724) | (0.0048) | (0.0205) | (0.0910) |
| PIPR | 0.8331 | 0.5780 | 0.4448 | 0.9005 | 0.6398 | 0.6183 |
|  | (0.0075) | (0.0324) | (0.0444) | (0.0259) | (0.0076) | (0.1023) |
| GNN-PPI | 0.8791 | 0.7472 | 0.6381 | 0.9226 | 0.8267 | 0.7137 |
|  | (0.0039) | (0.0526) | (0.0179) | (0.0010) | (0.0085) | (0.0533) |
| HIGH-PPI | 0.8797 | 0.7884 | 0.6749 | 0.8909 | 0.7567 | 0.6798 |
|  | (0.0089) | (0.0052) | (0.0211) | (0.0245) | (0.0103) | (0.0397) |
| SemiGNN-PPI | 0.8951 | 0.7832 | 0.7215 | 0.9240 | 0.8545 | 0.7178 |
|  | (0.0046) | (0.0315) | (0.0287) | (0.0022) | (0.0117) | (0.0356) |
| laruGL-PPI | 0.8762 | **0.7972** | **0.7407** | **0.9069** | **0.8061** | **0.7307** |
|  | (0.0053) | (0.0160) | (0.0672) | (0.0039) | (0.0096) | (0.0259) |
| HIGH-PPI* | 0.8480 | 0.7351 | 0.6665 | 0.8572 | 0.7127 | 0.6674 |
|  | (0.0067) | (0.0080) | (0.0452) | (0.0211) | (0.0277) | (0.0416) |
| laruGL-PPI* | **0.8635** | **0.7657** | **0.7097** | **0.9013** | **0.7852** | **0.7311** |
|  | (0.0048) | (0.0371) | (0.0394) | (0.0035) | (0.0110) | (0.0374) |

### 4.3 Performance on specific PPI types

For each of the six PPI types, we offer a separate performance analysis in terms of AUPR. It can be found that our presented model with lower learning cost parallels with HIGH-PPI as shown in Figure 4. Notably, laruGL-PPI consistently beats the previous state-of-the-art models in the most challenging setting (i.e. inductive setting with BFS). This is reasonable as laruGL-PPI learns spatial-biological patterns as possible as it can by sampling subgraphs in multiple graph frameworks. When facing unknown protein pairs, it can exploit these patterns to predict better than others.

**Figure 4.**
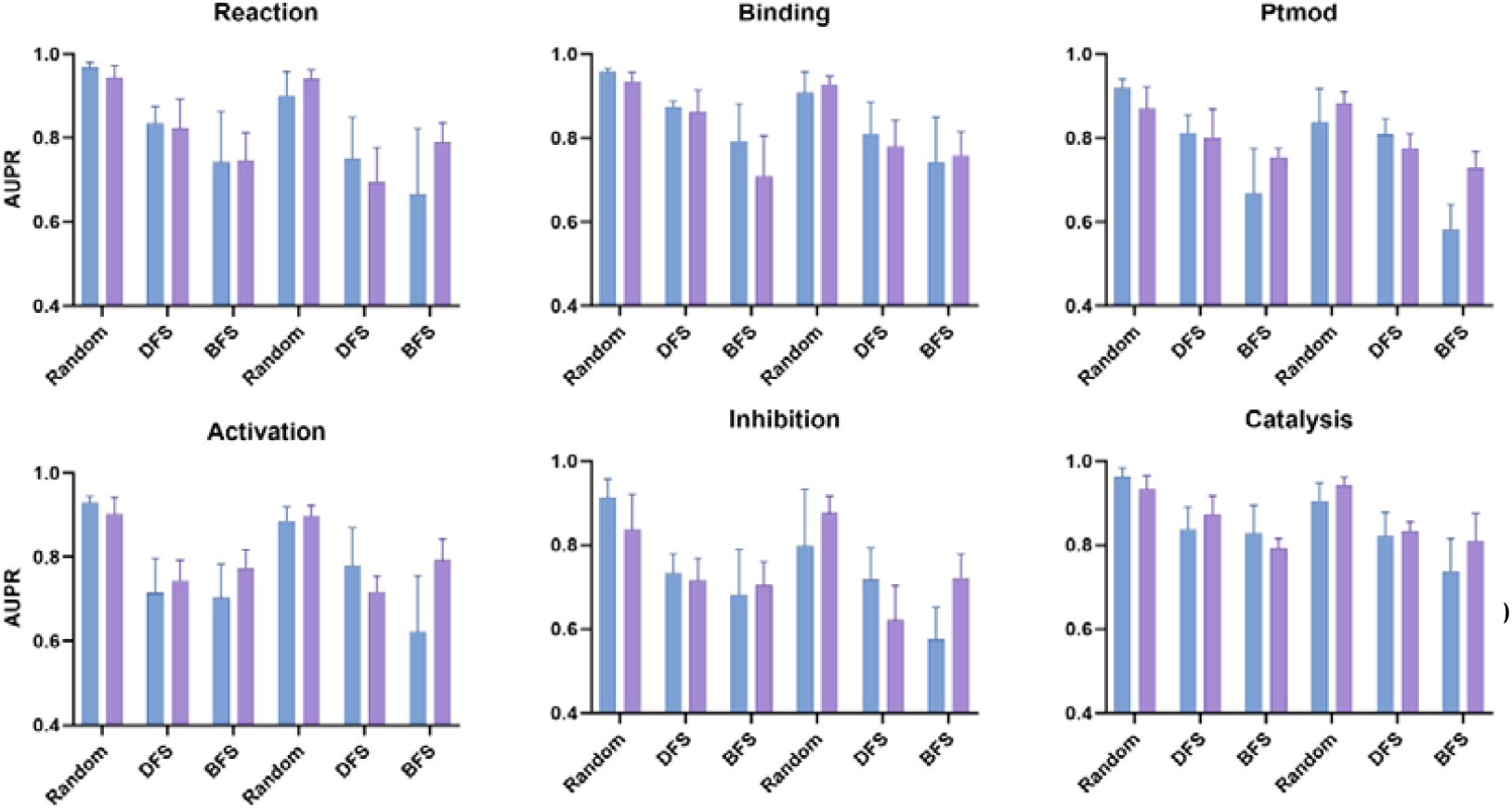
Performance of laruGL-PPI in predicting specific PPIs on SHS27k, compared with HIGH-PPI. Blue and purple represents HIGH-PPI and laruGL-PPI, respectively. Error bars express the standard deviation of the mean. Source data are provided as a Source Data file.

### 4.4 The results of training with different subgraph sizes

In this section, we display the results of the comparison of HIGH-PPI and our laruGL-PPI with various subgraph sizes. Beginning with a subgraph size of 200, we observe that our model has attained competing effectiveness in SHS27k under the transductive setting. From the results of the effect of subgraph size, using subgraph sampling is capable of reducing the memory cost of the Graphics Processing Unit (GPU) significantly and having a comparable outcome at the same time (Table S1). It is indicated that taking the size of 300, 400, or 500 can reach a greater performance under most scenarios (inductive setting with BFS segmentation) and request a relatively smaller memory cost in the SHS27k dataset (Figure 5). For the bigger dataset SHS148k, the advantages of models based on subgraph sampling are even more prominent. By sampling a subgraph of the size of 200, our model significantly surpasses the sota model called HIGH-PPI in difficult inductive scenarios with BFS. Specifically, our model achieves 0.7059 on the F1-score with 5.09 GB of computer graphics memory consumption in this condition, while the sota HIGH-PPI has a value of 0.6674, costing 28.93 GB (Figure 5). Generally, our model can significantly surpass the sota model in both transductive and inductive scenarios through suitable subgraph size (Table S2).

**Figure 5.**
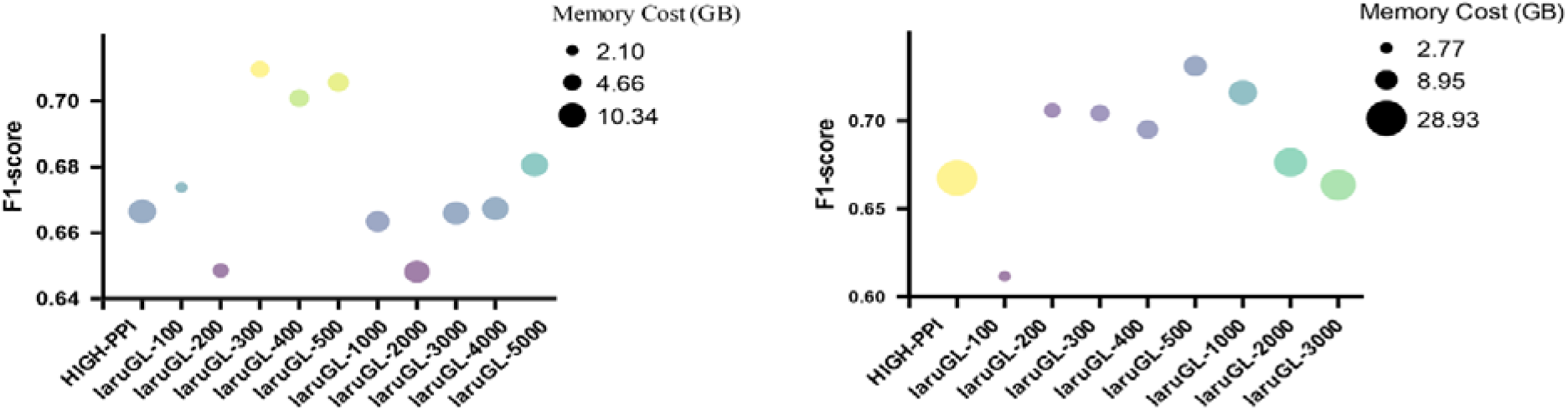
F1-score and Memory cost of HIGH-PPI and laruGL-PPI with various size subgraphs under inductive setting (BFS) on SHS27k(left) and SHS148k(right).

### 4.5 Ablation study

We investigate the effectiveness of different components in laruGL-PPI in Table 3. It can be seen that the model improves a lot (from 0.6749 to 0.7238) under relatively difficult circumstances (i.e. transductive setting with BFS) when considering adjacent residues in sequence. This implies that sequence information plays an important role in determining PPI types. What’s more, label graph encoding brings plenty of profits in a majority of scenarios, particularly under the most arduous condition (i.e. inductive setting with BFS, from 0.6652 to 0.7072). Finally, our model is getting more powerful when adopting an edge-based sampling strategy on the PPI graph. In particular, under a transductive setting, performance in terms of DFS achieves 0.7972. While under inductive setting, a more challenging scene, laruGL-PPI reaches the best F1-score in the light of all the partitions.

**Table 3.** Results of ablation studies on different components of laruGL-PPI using the SHS27k dataset.

| Model | Transductive setting |  |  | Inductive setting |  |  |
| --- | --- | --- | --- | --- | --- | --- |
|  | Random | DFS | BFS | Random | DFS | BFS |
| HGL-origin | 0.8797<br>(0.0089) | 0.7884<br>(0.0052) | 0.6749<br>(0.0211) | 0.8480<br>(0.0067) | 0.7351<br>(0.0080) | 0.6665<br>(0.0452) |
| HGL | 0.8873<br>(0.0059) | 0.7743<br>(0.0149) | 0.7238<br>(0.0383) | 0.8369<br>(0.0058) | 0.7400<br>(0.0165) | 0.6652<br>(0.0797) |
| laHGL | 0.8894<br>(0.0073) | 0.7819<br>(0.0226) | 0.7545<br>(0.0399) | 0.8405<br>(0.0040) | 0.7350<br>(0.0470) | 0.7072<br>(0.0543) |
| laruGL | 0.8762<br>(0.0053) | **0.7972**<br>**(0.0160)** | 0.7407<br>(0.0672) | **0.8635**<br>**(0.0048)** | **0.7657**<br>**(0.0371)** | **0.7097**<br>**(0.0394)** |

### 4.6 Interpretability of our laruGL-PPI

#### 4.6.1 Analysis of feature importance in residue-level

The feature importance in residue-level for overall (left-most column) and type-specific (right six columns) PPI prediction, which is calculated as the average z-score, resulting from dropping individual feature dimension from our model and calculating changes of AUPR before and after.

For node features in protein graphs, we select seven important features in terms of laruGL-PPI. Here, we list the selected seven residue-level physicochemical properties in Figure 5 and discuss their importance for different types of PPIs to both better interpret our model and discover enlightening biomarkers for the PPI interface. The average z-score, which results from deleting each feature dimension and analyzing changes in metrics before and after, is calculated to determine the importance of a feature. From the results, laruGL-PPI regards topological polar surface area (TPSA) and octanol-water partition coefficient (KOW) as dominant features in multiple types of PPIs. This finding supports the conventional wisdom that TPSA and KOW play a key role in the drug transport process[34], protein interface recognition[35, 36], and PPI prediction[37].

**Figure 5.**
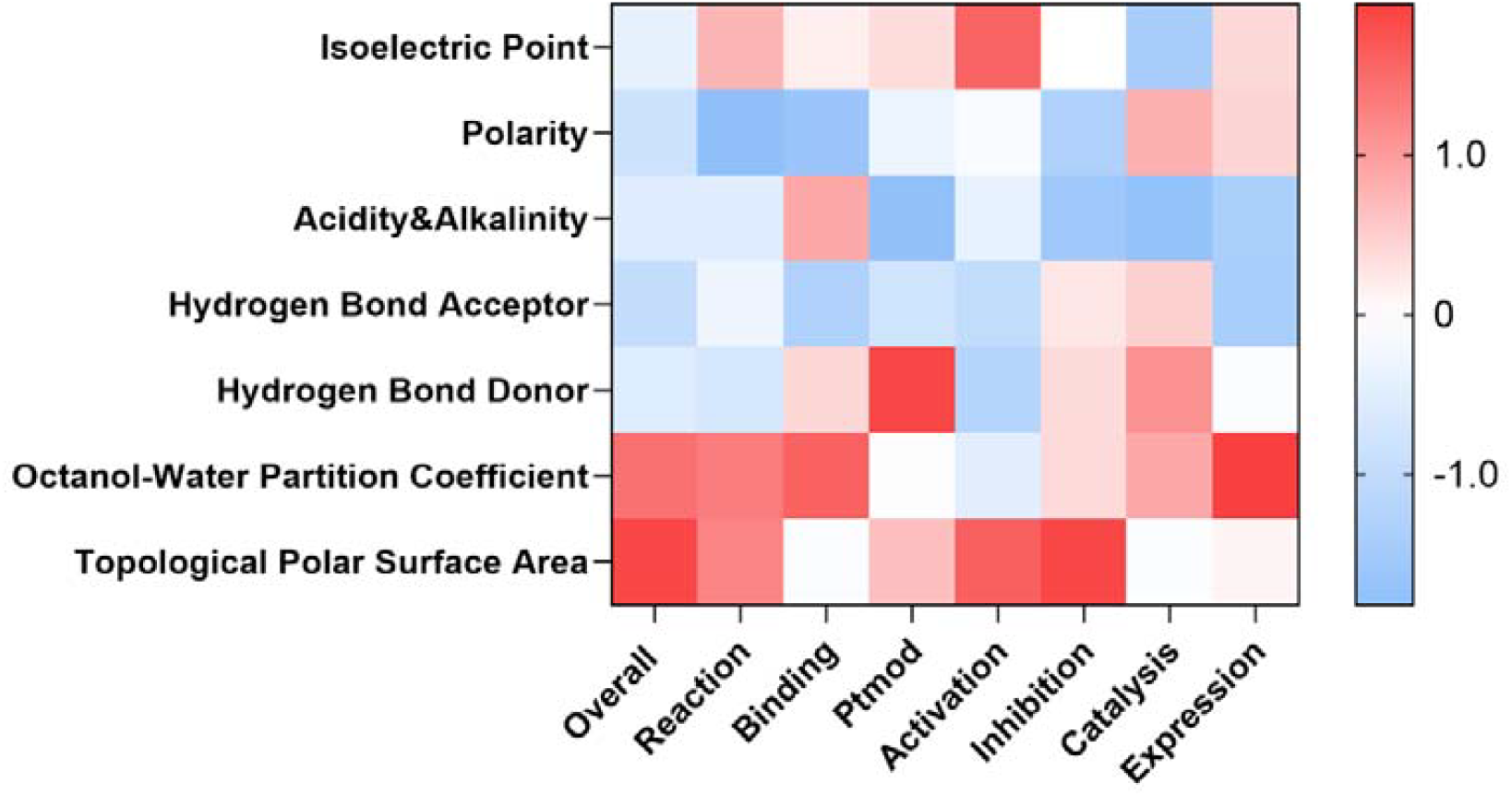
The feature importance in residue-level for overall (left-most column, F1-score) and type-specific (right seven columns, AUPR) PPI prediction.

#### 4.6.2 laruGL-PPI identifies key residues constituting functional sites

Typically, functional sites are spatially clustered sets of residues. They control protein functions and are thus important for PPI prediction. As our proposed model has the capacity to capture spatial-biological arrangements of residues in protein graph learning, this characteristic can be used to explain the model’s decision. It is meaningful to notice that laruGL-PPI can automatically learn the residue importance without any residue-level annotations. In this section, we provide a case study of predicting important residues for the binding surface and two cases of estimating key residues for catalytic function.

First, a binding example between the query protein (PDB id: 2B6H) and its partner (PDB id: 2REY) is investigated, whose complex structure is generated by ZDOCK[38]. Subsequently, we apply the GNN explanation approach on the laruGL-PPI model. As can be seen from Figure 6a. LaruGL-PPI can accurately and automatically identify the residues that belong to the binding domain.

**Figure 6.**
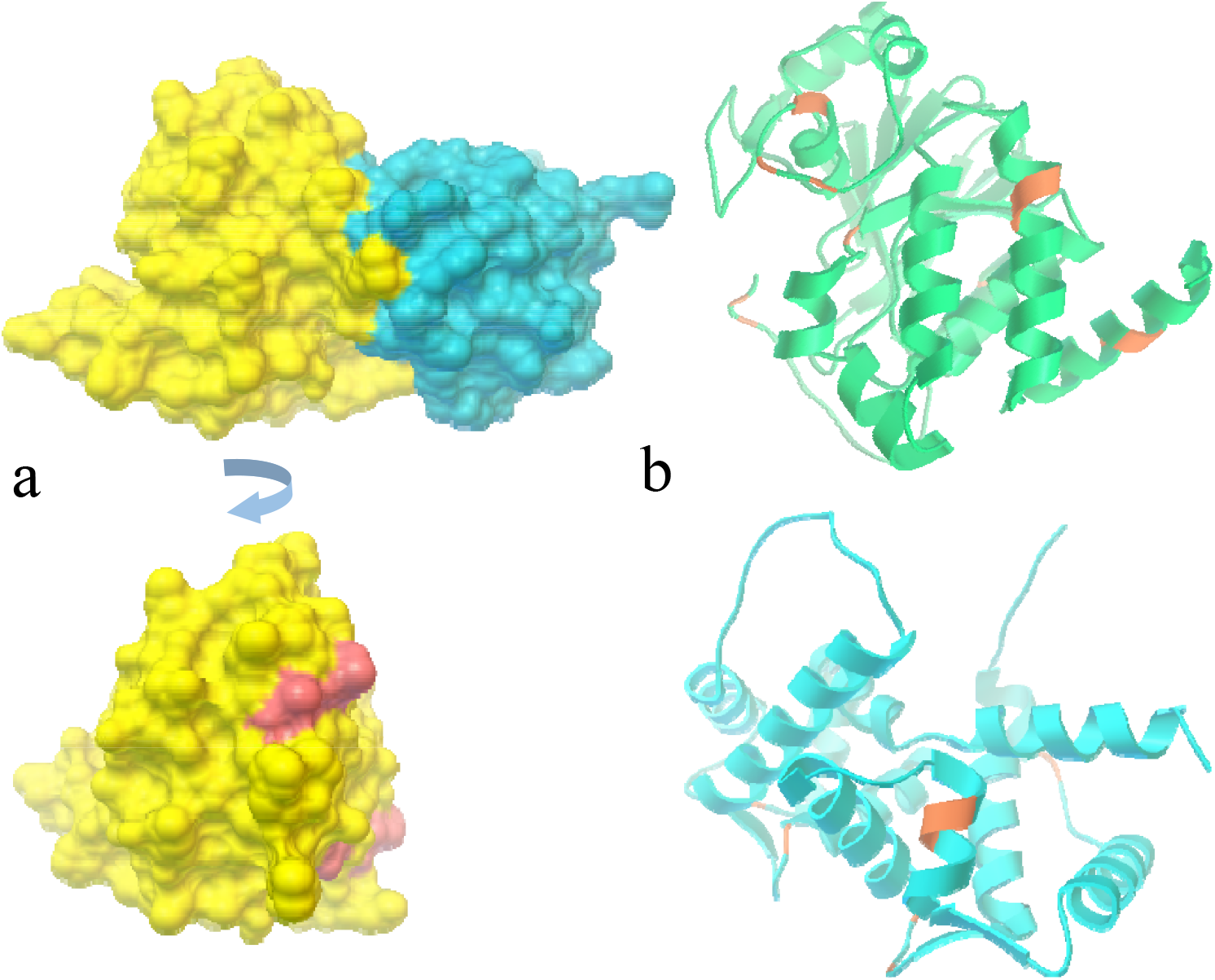
Automatic explanation for key residue without supervision. **a** Top: Depiction of a complex protein (left, query protein, PDB id: 2B6H; right, interacted protein, PDB id: 2REY) modeled in surface representation. Bottom: important residues of the query protein learned from laruGL-PPI with pink. **b** Depiction of two proteins (top, UniProt id: P62714; bottom, UniProt id: Q9UMX6) modeled in cartoon representations. Residues inferred by the model, which are colored with brown, are important for catalysis.

Second, to evaluate the prediction of crucial sites of catalysis for PPIs, we utilize the same GNN explanation approach in our model. The ground truth key sites, normally including binding sites and active sites, are retrieved from the Universal Protein Knowledgebase (UniProt)[39]. We explore the prominent residues of catalytic function for query proteins (UniProt id: P62714, Q9UMX6). As seen in Figure 6b, our proposed laruGL-PPI infers that ASN117 is important for the catalysis of P62714 known as the catalytic subunit of protein phosphatase 2A, in terms of Q9Y4X3. The coordinate bond between ASN117 and Mn^2+^ activates the enzyme to play a highly efficient catalytic role. For Guanylyl cyclase-activating protein 2 (Q9UMX6), the model can correctly predict that Asp66 and Asp102 are key to perform catalysis on Q16270.

Protein functional site prediction sheds light on the model decisions and how to conduct additional wet experiments for PPI investigation. Excellent model interpretability also demonstrates that our approach can accurately describe biological evidence for proteins.

## 5. Conclusions

In this study, it was found that conventional machine learning is subject to the problem of topological shortcuts. A new framework based on subgraph sampling was proposed to address this issue, enabling the model to focus more on the protein structure itself. This framework improved prediction accuracy and generalization, especially in challenging settings, where it exhibited significant improvements in encountering new proteins. Additionally, subgraph sampling saved computational resources. Through interpretable analysis of the model, it was found that the model can identify key sites, binding surfaces, or active sites. This may also be attributed to the subgraph sampling algorithm. Nevertheless, there is still room for improvement. For instance, the model does not explicitly consider protein sequence information, and future research may consider sequence-structure co-modeling. Another potential direction is to extend the model to protein all-atom graphs, extracting more microscopic information about proteins and learning the essence of PPI, thereby further enhancing the model’s generalization ability and interpretability.

## Data and code availability

The preprocessed data and model code associated with the study will then be uploaded to GitHub.

## CRediT authorship contribution statement

Yuanqing Zhou: Writing – review & editing, Writing – original draft, Visualization, Validation, Software, Methodology, Investigation, Conceptualization. Haitao Lin: Writing – review & editing, Software, Methodology, Conceptualization. Yufei Huang: Validation, Methodology, Investigation. Lirong Wu: Investigation, Conceptualization. Stan Z. Li: Supervision, Project administration, Funding acquisition. Wei Chen: Supervision, Project administration.

## Declaration of competing interest

None.

## Acknowledgments

This work was supported by the National Key Research and Development Program of China (No. 2022YFF1100202), Ministry of Science and Technology of the People’s Republic of China (No. 2021YFA1301603), National Natural Science Foundation of China Project (No. U21A20427), Project (No. WU2022A009) from the Center of Synthetic Biology and Integrated Bioengineering of Westlake University and Project (No. WU2023C019) from the Westlake University Industries of the Future Research Funding.

## Notes

### Competing Interest Statement

The authors have declared no competing interest.

### Summary of Updates

The revised drawings are more clear.

